# The Charcot-Marie-Tooth Neuropathy (CMTX3) Complex Structural Variation Causes Differential SOX3 Spatiotemporal Expression

**DOI:** 10.64898/2026.02.22.707254

**Authors:** Alexandra Boyling, Anthony N. Cutrupi, Desmond Li, Ben Crossett, Jonathan J. Danon, Edward D. Harvey-Latham, Garth A. Nicholson, Steve Vucic, Marina L. Kennerson

**Author notes:** Corresponding Author: Marina Kennerson, Northcott Neuroscience Laboratory, ANZAC Research Institute, Sydney Local Health District Gate 3 Hospital Rd, Concord, NSW, Australia 2139.

## Abstract

Charcot-Marie-Tooth (CMT) neuropathy is a clinically and genetically heterogeneous group of diseases characterised by the length dependent axonal degeneration of peripheral nerves. We previously mapped a rare form of X-linked CMT, CMTX3, to a 5.7-Mb interval on chromosome Xq26.3-q27.1 and excluded the coding region of all known genes in the linkage interval for mutations. Whole genome sequencing subsequently identified a 78-kb region of chromosome 8q24.3, that had been duplicated and inserted into the CMTX3 locus between the genes *HAPSTR2* and *SOX3*. The 78-kb insertion, which contains a partial transcript of *ARHGAP39*, fully segregated in families with CMTX3 and was absent in neurologically normal controls. To retain the CMTX3 insertion and investigate its consequences in appropriate neuronal tissue, we generated induced pluripotent stem cells (iPSC) from CMTX3 fibroblasts. Using bulk RNA sequencing of patient-derived spinal motor neurons, *ARHGAP39* was deemed non-pathogenic by excluding both the formation of novel fusion transcripts and dosage effects from the partial duplication. Subsequent NanoString expression analyses of candidate genes within the CMTX3 locus, across different stages of neuronal differentiation, identified spatiotemporal dysregulation of *SOX3.* NanoString showed reduced *SOX3* expression in patient iPSC. RNA sequencing detected *SOX3* downregulation in CMTX3 neuroepithelial progenitor cells, which was further confirmed by quantitative proteomics. Given the early onset and relatively rapid progression of CMTX3, these data prioritise *SOX3* as a leading candidate gene, consistent with its role as one of the earliest transcription factors expressed in the developing nervous system and a key regulator of neuronal fate.

## Introduction

Charcot-Marie-Tooth (CMT) neuropathy is a clinically and genetically heterogenous disease, causing length dependent axonal degeneration of motor and sensory neurons. Patients present with progressive distal muscle weakness and atrophy, sensory impairment, and variable involvement of the central nervous system. The disease has a prevalence of 1 in 2500^1^. CMTX3 is one of six X-linked forms of CMT^2^, with the most common being CMTX1, which is caused by disease associated variants in the gap junction beta 1 (*GJB1*) gene^3^. We mapped the CMTX3 locus in two distantly related families to a 5.7- Mb interval on chromosome Xq26.3-q27.3^4^. After excluding all genes within the linkage locus for coding mutations, selected affected individuals underwent short read whole genome sequencing. A complex structural variation (SV) was subsequently identified in which a 78-kb region of chromosome 8q24.3 was duplicated and inserted into the CMTX3 locus at a previously described Xq27.1 recombination “hotspot” between the genes *HAPSTR2* (formerly *LOC389895*) and *SOX3*^5^. Segregation analysis demonstrated the SV was present in all affected individuals (25 affected males and 30 carrier females) and absent in 50 unaffected family members as well as population controls^5^. This was the first report of a large insertion causing X-linked CMT and highlighted the importance of expanding the spectrum of mutation screening to structural variation in cases where all coding mutations had been excluded in a family.

Identification of the CMTX3 complex SV has contributed to the growing evidence that large DNA re-arrangements, particularly those located in non-coding or intergenic regions, may represent an under-recognised cause of inherited peripheral neuropathies. The change in the genomic landscape of the CMTX3 locus caused by the insertion introduced a duplicated portion of the *ARHGAP39* gene (NM_001308207.1; exons 1-7) from chromosome 8q24.3, with the closest genes to the 78-kb insertion being *HAPSTR2 (*formerly *LOC389895)* (located 329 kb proximal to insertion) and *SOX3* (located 84 kb distal to insertion). We proposed several hypotheses for the functional consequences of the CMTX3 insertion including gene dosage, the formation of novel fusion transcripts or alternative splicing, disrupting the interaction between a gene and its functional non-coding DNA sequences (such as promoter, enhancers and silencers), or introducing new regulatory interactions, resulting in dysregulated temporal and spatial gene expression^5^.

We have previously reviewed the impacts of SV in inherited peripheral neuropathies^6^ and devised a strategy for investigating the pathogenic consequences of the CMTX3 SV, particularly at the Xq27.1 “hotspot” where several other diseases have been reported due to the insertion of large, duplicated DNA fragments from different regions of the genome^7^. In the present study, we have used components of this strategy and generated patient-derived iPSC lines from re-programmed fibroblasts to retain the SV mutation and model CMTX3 at different stages of neuronal differentiation. We excluded the partial duplication of *ARHGAP39* as a causative mechanism and, using complementary transcriptomic approaches together with proteomics, identified *SOX3* as the lead candidate gene for CMTX3 based on its spatiotemporally specific downregulation in CMTX3-derived tissue.

## Materials and Methods

### Ethics statement

The protocols utilised in the current study (2019/ETH07839) have been approved by the Sydney Local Health District Human Ethics Review Committee. Informed consent was obtained from study participants.

### Cell Line Summary

Although CMT is classified as a rare disorder, CMTX3 is ultra-rare because of its exceptionally low prevalence^8^, creating clear constraints on access to sufficient donor material for functional studies. Accordingly, the sample sizes in this study range from n=1 to n=3 CMTX3-derived cell lines, reflecting the staged acquisition and expansion of donor material over the course of the project. Full details of the cell lines used in the present study are provided in Supplementary Table 1.

### Culturing and maintenance of primary fibroblasts

Primary fibroblasts were maintained in F-DMEM culture medium comprising DMEM, 10% (v/v) Foetal Bovine Serum, 1% (v/v) Penicillin/Streptomycin and 1% (v/v) L-glutamine (Gibco, Thermo Fisher Scientific, Waltham, MA, USA). All cells were maintained in 5% CO_2_, humidified air at 37°C. Media was replaced every 2-3 days. Cells were maintained under these conditions for 7 days for biobanking or harvested (70-80% confluency) for downstream experiments.

### Generation, culturing, and maintenance of iPSC lines

Reprogramming of patient and unrelated, neurologically normal control fibroblasts was outsourced to the Stem Cell and Organoid Facility (SCOF; Children’s Medical Research Institute, Westmead, NSW, Australia) and FUJIFILM Cellular Dynamics (Madison, WI, USA), where in-house immunofluorescence was used to confirm the expression of multiple pluripotency markers. Reprogrammed iPSC were shown to be karyotypically normal using either G-banded karyotyping (WiCell; Madison, WI, USA) or digital SNP karyotyping with the Illumina Infinium GSA-24 v3.0 microarray (Victorian Clinical Genetics Services, Melbourne, VIC, Australia). Description of reprograming methods used by FUJIFILM Cellular Dynamics have been detailed elsewhere^9^. Additional control lines were purchased from FUJIFLM Cellular Dynamics. iPSC were seeded in 6-well plates (Corning, New York, USA) coated with Matrigel (0.167 mg/mL, Corning) and maintained in TeSR-E8 medium (STEMCELL Technologies, Vancouver, BC, Canada) that was replaced daily. Subculturing was performed using 0.5 mM UltraPure EDTA (Invitrogen, Carlsbad, CA, USA) as described in Cutrupi et al. (2023)^9^.

### Differentiation of neuronal precursors and mature spinal motor neurons

iPSC were differentiated into caudal neuroepithelial cells (NEP – differentiation day 6), motor neuron progenitors (MNP – differentiation day 12), neurospheres (NSPHR – differentiation day 18) and mature spinal motor neurons (sMN – differentiation day 28) using previously described methods^9^. SN38-P was used to further purify neuronal cultures from non-neuronal cells^10^, as described in^9^. Adherent culturing was performed in 6-well plates (Corning-Costar) pre-treated with 0.167 mg/mL Matrigel (Corning). Suspension culturing was performed in ultra-low attachment 6-well plates (Corning-Costar).

### CMTX3 SV genotyping PCR assay

DNA was isolated from patient and control iPSC using QuickExtract DNA Extraction Solution 1.0 (Epicentre, Madison, WI, USA) according to the manufacturer’s instructions. A multiplex PCR was used to genotype the CMTX3 complex insertion as previously described^5^, with primers synthesised by Sigma Aldrich (St. Louis, MO, USA). Agarose gels (2% w/v) were prepared in 1 X TAE buffer (Astral Scientific) with 0.01% SYBR Safe DNA gel stain (Thermo Fisher Scientific). HyperLadder 100 bp (Bioline, London, UK) was used as a size-standard. PCR products were size fractionated and visualised using a Safe Imager Transilluminator 2.0 (Invitrogen). Images were captured using a Canon PhotoShot S5-IS digital camera with Hoya O(G) filter.

### Quantitative real-time polymerase chain reaction (qRT-PCR) for the confirmation of iPSC pluripotency

Total RNA was isolated from CMTX3 (P1, P2) and control (C1-C3) iPSC as well as a single control fibroblast line (negative control) using the RNeasy Mini Kit (Qiagen, Hilden, Germany). Reverse transcribed template was prepared using the iScript cDNA Synthesis Kit (BioRad, Hercules, CA, USA). Quantitative RT-PCR was performed using Taqman Gene Expression Assays for 6 pluripotency markers (*LIN28A, NANOG, NODAL, FOXD3, CDH1, TDGF1*) and one internal housekeeping gene (*GAPDH*) (Supplementary Table 2). Reactions were prepared in 20 µL volumes containing 1X TaqMan Gene Expression Assay (Applied Biosystems, Foster City, CA, USA), 1X TaqMan Gene Expression Mastermix (Applied Biosystems) and 25ng cDNA template. Water was used as a negative control in all assays. Thermal cycling was performed (StepOnePlus Real-Time PCR; Applied Biosystems) using the following cycling protocol: 95°C for 20 s, followed by 40 cycles of 95°C for 1 s and 60°C for 20 s. Data analysis was performed using the StepOne Plus Software (version 2.1). GraphPad Prism (version 10) was used to plot the raw Ct values, enabling a qualitative representation of the expression of pluripotency genes between iPSC and fibroblasts^11,12^.

### NanoString nCounter targeted gene expression analysis

Gene expression quantification using the NanoString nCounter gene expression system (NanoString Technologies, Seattle, WA, USA) was outsourced to Westmead Medical Research Institute (Sydney, NSW, Australia). A custom NanoString nCounter Elements TagSet panel was designed to detect 48 RNA targets including 21 genes within and flanking the CMTX3 linkage region, 1 gene contained within the CMTX3 insertion sequence, 11 genes implicated in peripheral nervous system differentiation and 14 housekeeping genes (Supplementary Table 3). Housekeeping genes were selected in accordance with the NanoString technical guidelines which recommend utilising a combination of housekeeping genes with high, medium and low expression levels^13^. The target-specific oligonucleotides were designed by NanoString Technologies (Seattle, WA, USA) and synthesized by Integrated DNA Technologies (Coralville, IA, USA). Total RNA was isolated from patient (n = 2) and control (n = 3) tissues using the RNeasy Mini Kit (Qiagen) and diluted to a concentration of 14.286 ng/µL. RNA yield was assessed using NanoDrop (Thermo Fisher Scientific) and integrity determined using a TapeStation (Agilent Technologies, Santa Clara, CA, USA). Details regarding assay preparation, execution and analysis are previously described^9^. For analyses involving multiple different cell types where housekeeping expression levels were quite variable, a selection of housekeeping genes showing minimal variability was utilised. The nSolver Analysis Software (NanoString Technologies) was used for statistical analysis of differential gene expression between patient and control samples. A Welch’s t-test was performed using the log2-transformed normalised count data as input and a significance threshold of p < 0.05.

### RNA sequencing data processing and analysis

Total RNA was extracted from sMN cells (n = 1 patient (2 clones), n = 3 controls) and NEP cells (n = 2 patients, n = 3 controls) using the RNeasy Mini Kit (Qiagen). Yields were quantitated using NanoDrop (Thermo Fisher Scientific) and the RNA integrity determined using a 2200 TapeStation (Agilent Technologies). Poly(A) RNA-Seq, including library preparation and data processing was outsourced to Macrogen (Seoul, South Korea) as previously described^9^. Briefly, novel fusion transcripts were predicted using three bioinformatic tools; DeFuse^14^, FusionCatcher^15^ and Arriba^16^. To identify novel transcript and splice variants, transcripts were assembled using StringTie^17^ and then classified with GffCompare^18^. Gene-level differential gene expression analysis was performed using edgeR^19^ and/or DESeq2^20^. Differentially expressed genes (DEGs) were determined by log_2_FC *≠* 0 and a false discovery rate (FDR) adjusted *p-*value threshold of < 0.05 (edgeR) or < 0.1 (DeSeq2).

### Immunofluorescence

Immunofluorescence experiments were performed as previously described^9^. In brief, cells were cultured in Nunc Lab-Tek 8-well chamber slides (Thermo Fisher Scientific) or black, optically clear PhenoPlate96-well microplates (Perkin Elmer, Waltham, MA, USA) coated in Matrigel (0.167 mg/mL; Corning). Cells were washed once in DPBS (Gibco), fixed in 4% (v/v) paraformaldehyde (PFA, Sigma Aldrich) for 20 min at room temperature, washed once with DPBS, treated with permeablisation solution (DPBST) containing DPBS and Triton X-100 (Calbiochem) ranging from 0.2-0.3% (v/v) (depending on density of cellular clumps) for 30 min at room temperature and blocked (DPBS and 5% (w/v) bovine serum albumin (Sigma Aldrich)) for 1 h at room temperature. Cells were incubated with primary antibodies overnight at 4°C. iPSC: anti-OCT4 (Cell Signaling Technologies, Danvers, MA, USA, #2840, 1:400), anti-SOX2 (Cell Signaling Technologies, #3579, 1:400), and anti-NANOG (Cell Signaling Technologies, #4903, 1:400). MNP: anti-OLIG2 (Millipore, Burlington, MA, USA, MABN50 1:100). sMN: anti-MNX1 (Sigma, HPA071717, 1:500), and anti-TUBB3 (Sigma, T2220, 1:1000). The cells were then incubated with Alexa Fluor (AF) secondary antibodies (Invitrogen) for 1 h at room temperature. The following secondary antibodies were used: goat anti-rabbit AF 488 (Invitrogen, A11008, 1:500) goat anti-rabbit AF 555 (Invitrogen, A21428, 1:500), goat anti-mouse AF 488 (Invitrogen, A11001, 1:500), and goat anti-mouse AF 555 (Invitrogen, A21422, 1:500). Nuclei were counter-stained with 4,6-diamidino-2- phenylindole (DAPI, Molecular Probes, 0.2 µg/mL) and mounted with mounting media (75% glycerol (v/v) and 20 mM Tris, pH 8). Samples were imaged on a Leica SP8 confocal microscope (Leica Microsystems, Wetzlar, Germany) and visualised using Leica Application Suite X (LAS X) software (Leica Microsystems). Images were processed using FIJI^21^ (version 2.14.0) for Windows.

### Quantification of MNX1 positive sMN

FIJI (version 2.14.0) was used to generate maximum intensity Z-projection images for downstream analysis. Images of sMN were obtained from 3 separate rounds of differentiation (n = 2 patients, n = 3 controls), with cells stained 8 days after neuron plating. A custom CellProfiler^22^ (version 4.2.5) pipeline was used to calculate the percentage of sMN nuclei positive for MNX1. First, the ‘IdentifyPrimaryObjects’ module was used to identify DAPI-stained nuclei. A global minimum cross-entropy threshold method was employed. To determine the lower threshold limit, images were manually inspected using the ‘pixel intensity’ tool, to provide intensity values for background fluorescence. To determine the nuclei diameter range, the ’measure length’ tool was used to manually assess the typical diameter of nuclei within the fluorescence images. Optimal segmentation of nuclei was achieved by de-clumping objects based upon intensity. However, the presence of small, bright DAPI speckles introduced the risk of false-positive nuclei identification, which could have significant effects on downstream analysis. To overcome this issue, the ‘FindMaxima’ module was used to identify high-intensity DAPI speckles, which were then processed using the ‘ConvertImageToObjects’ module and subsequently excluded from the image using the ‘MaskObjects’ function. Masking settings were optimised to retain partially masked objects if they display high overlap with the masking region. This setting minimises the risk of nuclei being discarded in cases where they co-localise with the DAPI speckles. To identify the number of MNX1-positive cells, the MNX1-stained images were processed using the ‘IdentifyPrimaryObjects’ module, as described above. The ‘RelateObjects’ module was used to associate the two sets of identified objects (i.e. the DAPI objects and the MNX1 objects) with each other. Nuclei were then filtered based on the presence of MNX1-staining using the ‘FilterObjects’ module. The number of MNX1- positive nuclei was divided by the total number of nuclei to calculate the proportion of sMN showing positive expression of MNX1. Proportion data underwent arcsine square root transformation prior to statistical testing. Statistical analysis was performed in GraphPad Prism (version 10) using an unpaired two-tailed t-test with a significance threshold of p < 0.05.

### Quantification of SOX3 by tandem LC-MS/MS

Cell pellets were resuspended in lysis buffer (4% sodium deoxycholate, 100 mM Tris, pH 8.5) and heated at 95 °C for 10 min. Pellets were lysed by sonication using a Branson Digital Sonifier SFX 150 (Emerson, MO, USA) using 2 rounds of 30% power for 30 s each. Debris was pelleted by centrifugation at 16,000 rcf for 10 min and supernatant collected. Cysteines were reduced and alkylated by incubation with 10 mM Tris(2-carboxyethyl)phosphine (TCEP) and 40 mM chloroacetamide at 95 °C for 10 min. Samples were diluted 4-fold with water and digested with sequencing grade modified porcine trypsin added in a 1:20 ratio (trypsin:protein content) for 16 h at 37 °C. Digested samples were cleaned by SDB-RPS stage tips (styrenedivinylbenzene - reverse phase sulfonate) as per^23^. Briefly, samples were mixed with an equal volume of 99% ethyl acetate, 1% TFA to extract the SDC and centrifuged. The lower phase was cleaned by binding to a SDB-RPS stage tip and washed with 99% ethyl acetate,1% TFA then 0.2% TFA and eluted using 5% ammonium hydroxide, 80% acetonitrile and dried. Samples were reconstituted in 50 mM TEAB (triethylammonium bicarbonate), pH 8.5 and 20 µg of peptide from each sample labelled with TMT6plex isobaric labelling reagent (Thermo Fisher Scientific), following the manufacturer’s instructions. The TMT labelled peptides were combined and fractionated by high pH reverse phase chromatography using a 4.6 mm x 250 mm C18 column (3.5 μm particle size, Xbridge Premier BEH C18 ; Waters, MA, USA) on a Vanquish fraction collector (Thermo Fisher Scientific) with mobile phases A (10 mM ammonium formate, 2% v/v acetonitrile, pH 8.5) and B (10 mM ammonium formate, 80% v/v acetonitrile). Peptides were eluted using a linear gradient of 10% to 40% B over 11 mins at a flow rate of 1 mL/min. 16 concatenated fractions were collected and dried down prior to LC-MS analysis.

Fractions were reconstituted in 3% acetonitrile v/v, 0.1% formic acid v/v and separated on an in-house packed column (75 μm x 500 mm, 1.9 μm ReproSil Pur 120 C18, Dr. Maisch GmbH, Germany) with a Vanquish Neo LC (Thermo Fisher Scientific) with mobile phase A (0.1% v/v formic acid) and B (80% v/v ACN, 0.1% FA v/v). Peptides were eluted at a flowrate of 0.3 μL / min and a gradient of 3% B to 40% B in 66 min, 40% B to 60% B in 2 min, 60% B to 98% B in 2 min, and then washed with 98% B whilst increasing flow from 0.3 to 0.5 μL / min over 5 min. The column was then equilibrated at 3% B for 2.7 column volumes at 750 bar. The LC was coupled to an Orbitrap Eclipse mass spectrometer (Thermo Fisher Scientific) with a spray voltage of 2300 V, RF lens of 30%, and ion transfer tube heated to 300 °C. The Orbitrap Eclipse was operated in data-dependent acquisition (DDA) mode comprising of a survey MS scan acquired as profile with maximum injection time of 118 ms and standard AGC target across a scan range from 350 – 1400 m/z with Orbitrap resolution of 60,000 with a 1.5 s cycle time. MS/MS scans were performed between MS scans isolating the most abundant ions (1.4 m/z isolation window, >5e4 ions) with charge states +2 to +4 and a dynamic exclusion of 30 s. MS/MS scans were acquired with a maximum injection time of 54 ms and a normalised AGC target of 400% with a scan range starting from 100 m/z, HCD collision energy of 35% and Orbitrap resolution of 30,000.

The data was analysed using Proteome Discoverer vr 2.5 (Thermo Fisher Scientific) and Mascot vr 3.1^24^ (Matrix Science, London, UK) using machine learning model MS2PIP:HCD2019^25^. The search parameters included the following variable modifications: acetylation (protein N-terminal), oxidation (M), deamination (NQ), carbamidomethyl (C), TMT6plex (K and N-term) and trypsin enzyme allowing for 2 missed cleavages and 10 ppm precursor and fragment mass tolerance. The Swiss-Prot Homo Sapiens database (<u>UP000005640</u>) was used and results filtered at 1% FDR (false discovery rate). The TMT reporter fragments were used for quantification. Differential protein expression was performed on RStudio using the Differential Expression analysis of Proteomics data (DEP) tool^26^.

## Results

### Characterisation of CMTX3 patient-derived iPSC lines

Pluripotency of CMTX3 iPSC was confirmed by FUJIFILM Cellular Dynamics Incorporated and SCOF using in-house protocols. Upon receipt of the patient-derived iPSC, re-confirmation of pluripotency marker expression was performed at both the RNA and protein level using qPCR and immunofluorescence respectively (Figure 1A-B). To ensure the CMTX3 SV was preserved following cellular reprogramming, patients and controls were genotyped by the multiplex PCR previously designed to detect the CMTX3 SV breakpoints^5^. Figure 1C-D shows confirmation the CMTX3 breakpoint junction fragments are amplified in the patient and absent in control iPSC lines.

**Figure 1:**
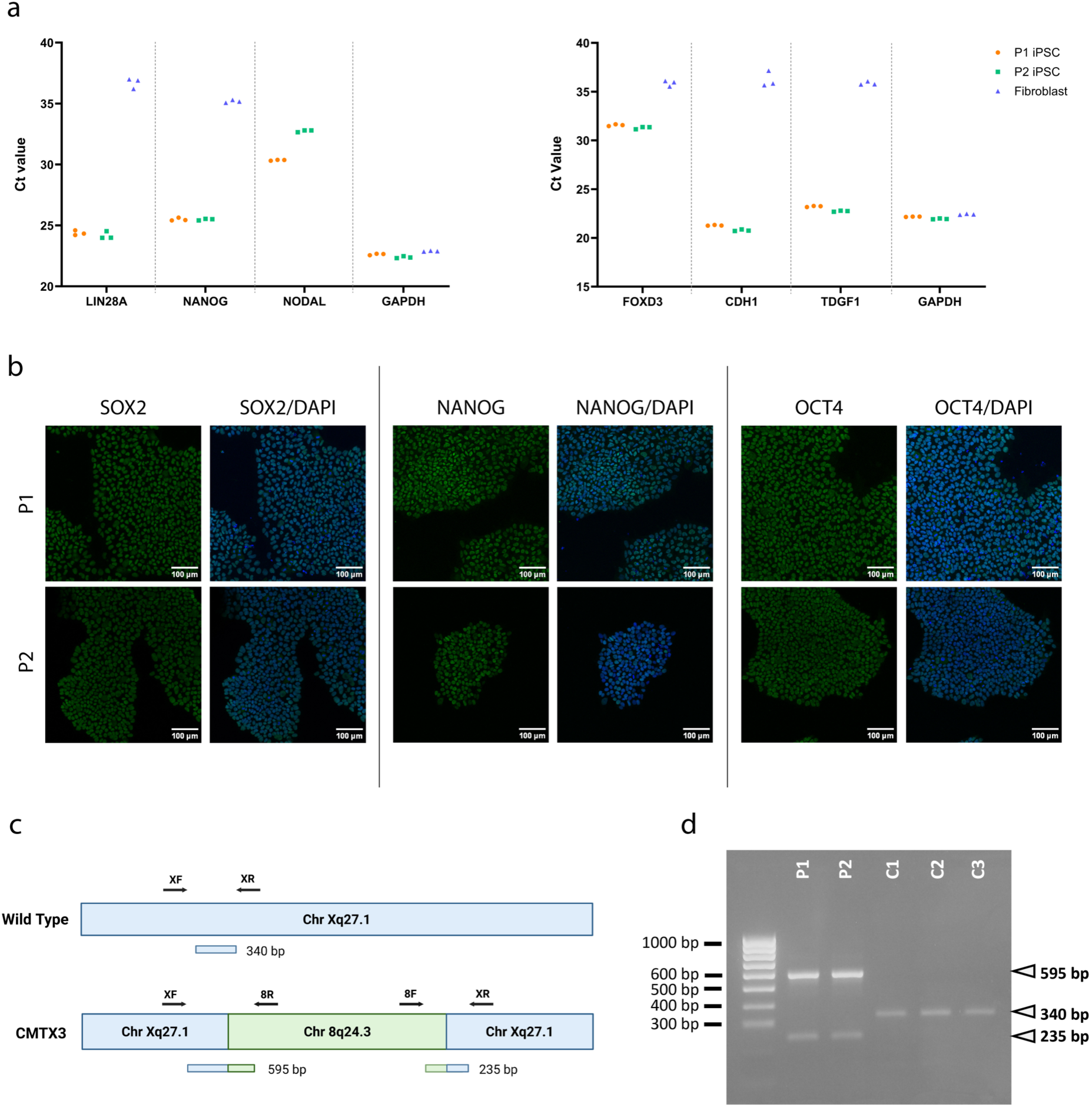
Characterisation and genotyping of CMTX3 patient-derived iPSC. (A) RT-qPCR assessment of pluripotency-associated transcripts (LIN28A, NANOG, NODAL, FOXD3, CDH1, TDGF1) in two independent CMTX3 iPSC lines (P1 and P2) compared to fibroblasts. GAPDH was used as an endogenous control. As NODAL was undetectable in fibroblasts, data is represented as raw Ct values. (B) Immunofluorescence staining confirming pluripotency marker expression, (SOX2, NANOG and OCT4; green) in CMTX3 iPSC (P1 and P2). Nuclei are counterstained with DAPI (blue). Scale bars, 100 µm (C) Schematic of the CMTX3 structural variant and genotyping primer (F = forward, R = reverse) locations indicating the expected amplicons from the wild type X chromosome (340 bp) and CMTX3 junction fragments (595 bp and 235 bp) (adapted from Brewer et al.6, created with BioRender). (D) Genotyping PCR and gel electrophoresis confirms the presence of the CMTX3 SV in patient-derived iPSC lines (P1 and P2), and their absence in control lines (C1-C3). HyperLadder 100 bp (Bioline) was used as a size standard. Agarose gel (2% w/v) was prepared in 1X TAE buffer containing 0.01% SYBR Safe (Bioline).

### Generation of MNP and sMN by differentiation of CMTX3 patient-derived iPSC lines

To generate a motor neuron model for CMTX3, 2 patient and 3 control iPSC lines were differentiated into sMN from highly expandable *OLIG2*-positive motor neuron progenitors (MNP) as previously described^9^. Immunofluorescence confirmed efficient ventralisation with differentiating cells expressing OLIG2 at the MNP stage (Supplementary Figure 1A). NanoString nCounter profiling consistently showed *OLIG2* expression was higher in MNPs than in iPSC, where expression appears undetectable (Supplementary Figure 1B), supporting successful derivation of MNPs. Terminal differentiation of MNP to mature sMN was performed under 3D culture conditions as described previously^9^. Mature sMN showed robust expression of the sMN marker MNX1 and the neuronal cytoskeletal marker βIII-Tubulin (TUBB3) (Figure 2A). The proportion of MNX1-postive sMN did not differ between CMTX3 patients (76.53% ± 3.966 SEM) and controls (69.99% ± 1.418 SEM) (Figure 2B; *p* > 0.05). Likewise, NanoString analysis demonstrated increased expression of the sMN markers *MNX1, ISL1* and *ChAT*, in sMN relative to iPSCs and MNPs (Figure 2C (i)) with no significant differences between CMTX3 and control lines (Figure 2C (ii)); *p* > 0.05).

**Figure 2:**
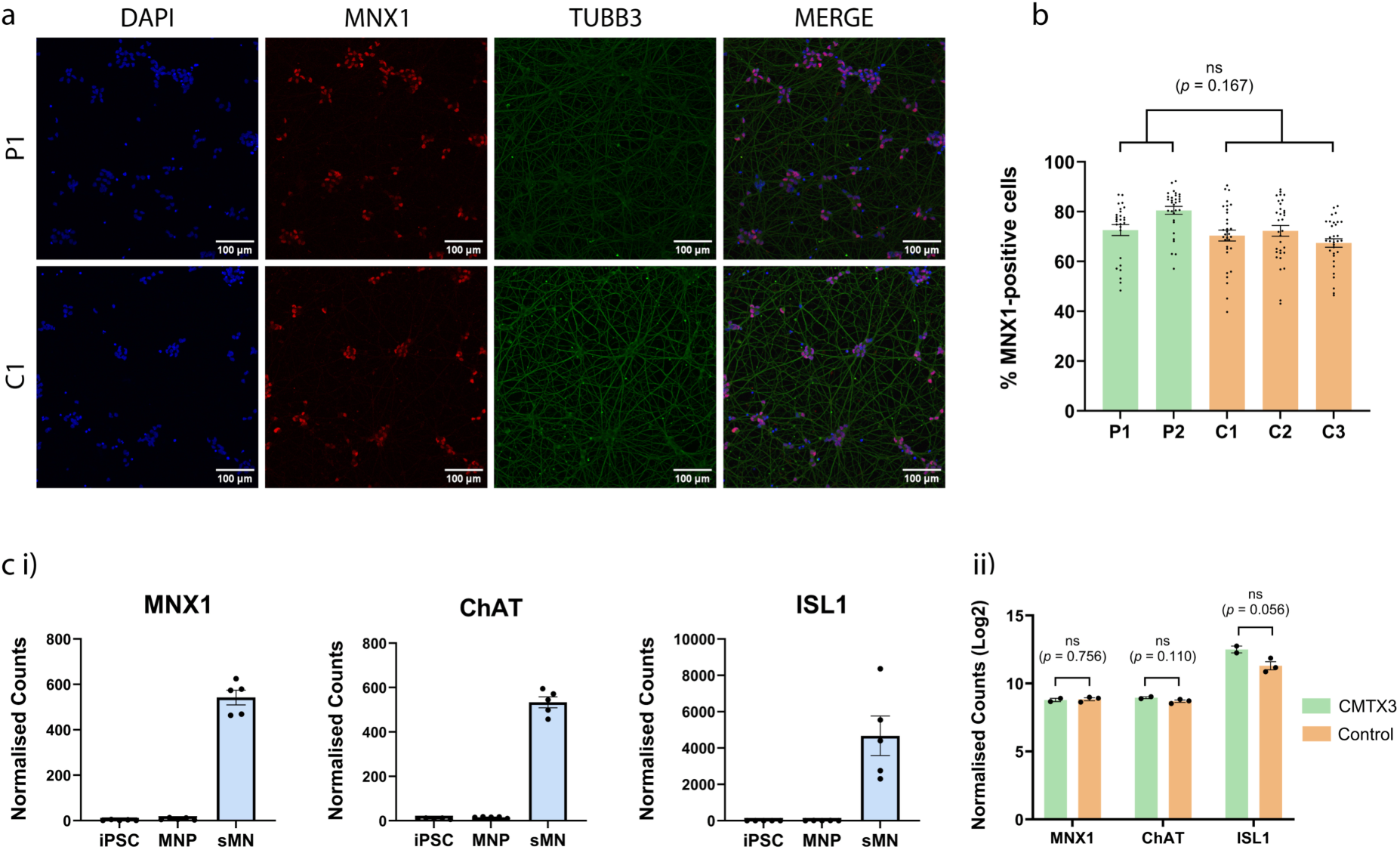
iPSC-derived sMN express canonical motor neuron markers. (A) Representative immunofluorescence images of terminally differentiated sMN from a CMTX3 patient (P1) and control (C1) showing MNX1 (motor neuron identity; red) and βIII-tubulin/TUBB3 (neuronal cytoskeleton; green). Nuclei are counterstained with DAPI (blue); merged images are shown. Scale bars, 100 μm. (B) Quantification of MNX1-positive nuclei across CMTX3 patient (P1 and P2) and control (C1-C3) lines. Points represent the proportion of MNX1-positive nuclei from acquired images collected over 3 independent differentiations. Bars represent mean ± SEM. No significant difference in MNX1+ yield was detected between CMTX3 and controls (unpaired two-tailed Student’s t-test; ns p=0.167). (C) NanoString nCounter analysis of motor neuron marker expression (i) Normalised mRNA counts for MNX1, ChAT, ISL1 are shown at various differentiation stages (iPSC, MNP, sMN; n=5 cell lines per stage). (ii) Log2-transformed normalised counts for the same markers in sMN, comparing CMTX3 patient (n=2) versus control (n=3) samples. Statistical analysis: unpaired t-test with Welch’s correction and a significance threshold of p < 0.05. ns: non-significant

### Exploratory RNA-Seq of patient-derived sMN reveals no novel fusion transcripts or aberrant splicing associated with the CMTX3 structural variant

Two leading hypotheses have been proposed to explain the pathomechanism of CMTX3. The first centres on the partial *ARHGAP39* sequence contained within the CMTX3 insertion, suggesting disease could arise from overexpression of a truncated *ARGHAP39* transcript or the generation of a novel transcript species^5^. The second proposes that the ∼78 kb SV perturbs the regulatory landscape, leading to dysregulation of one or more nearby genes^5^. RNA-Seq allows simultaneous interrogation of both hypotheses in a single experiment and was therefore used to provide an initial evaluation of the transcriptional consequences of the CMTX3 SV in sMN, using patient-derived (n=1 individual, two iPSC clones) and controls (n=3 lines). Quality control metrics are available in Supplementary Table 4.

None of the three fusion-calling tools identified fusion transcripts with breakpoints spanning the CMTX3 linkage region on chromosome X and the duplicated 8q24.3 fragment (Supplementary Table 5). Among fusion transcripts with both breakpoints on chromosome X, none mapped to the CMTX3 linkage interval (Supplementary Table 6-8). Although deFuse reported putative *ARHGAP39*-associated fusions, these were not considered as pathogenic because (a) one of the two breakpoints lay outside the CMTX3 insertion sequence, and (b) identical transcripts were also predicted in all three control lines (Supplementary Table 9). No additional fusion transcripts involving the duplicated fragment of 8q24.3 were detected (Supplementary Tables 10-11).

No predicted novel transcripts incorporated sequence from either the CMTX3 insertion fragment (Supplementary Table 12A) or the broader CMTX3 locus at Xq27.1 (Supplementary Table 12B). Within the CMTX3 linkage interval, eight novel splice variants were detected, however none were specific to patient-derived samples (Supplementary Table 13A). There were no novel splice variants originating from within the 8q24.3 insertion sequence (Supplementary Table 13B).

To investigate if CMTX3 involves local gene dysregulation, differential gene expression analysis was performed. Because biological replicates for CMTX3 were not available in this preliminary experiment, the analysis was exploratory and intended to nominate candidate genes for further follow-up. Differential expression was analysed using edgeR which can be applied to datasets with limited sample sizes^19^. In total, 51 differentially expressed genes were predicted. None mapped to either the CMTX3 linkage interval at Xq27.1 or to the duplicated ∼78 kb 8q24.3 fragment (Supplementary Table 14). To specifically evaluate local effects, Table 1 summarises log2(fold change) values for positional candidate genes within the CMTX3 locus, providing an overview of local gene regulation, although none reached statistical significance (FDR > 0.05). Of the 29 positional candidates, 17 were filtered by edgeR due to low expression in sMN. Remaining genes showed largely modest changes with average log(fold change) < 0.5 (Table 1). Two genes, *SOX3* and *ZIC3*, showed larger magnitude shifts, however, *ZIC3* expression was highly discordant between patient clones (clone A= −2.185, clone B= −0.094) (Table 1). In contrast, *SOX3* showed a consistent trend of down-regulation across both clones (Clone A logFC −1.383 and Clone B logFC −1.341), albeit at low overall abundance (mean TPM 0.567 in the patient clones and 1.477 TPM in controls).

**Table 1:**
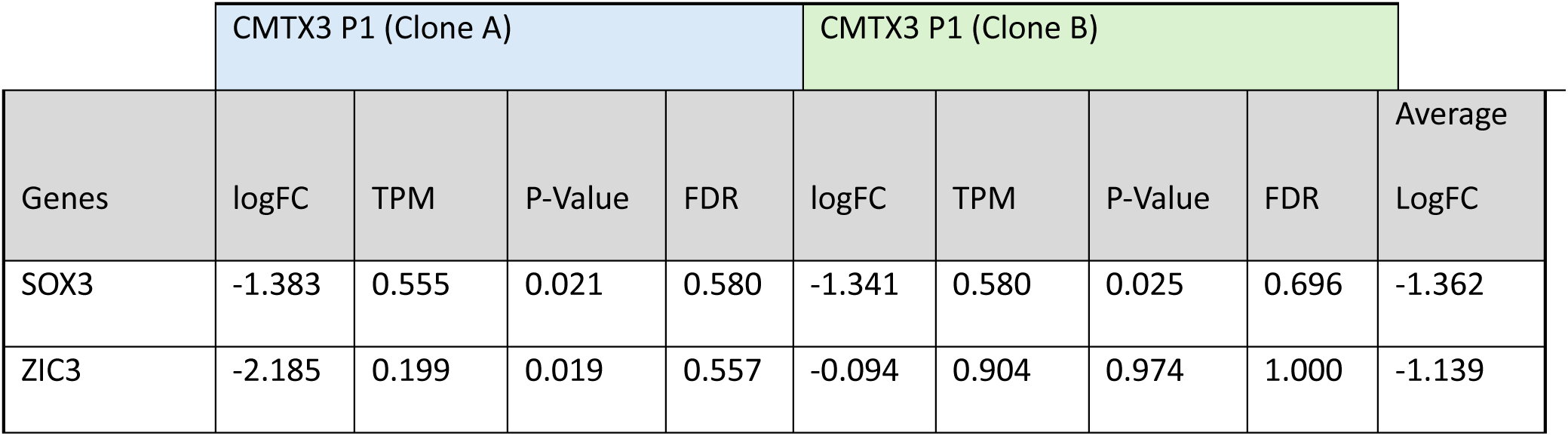
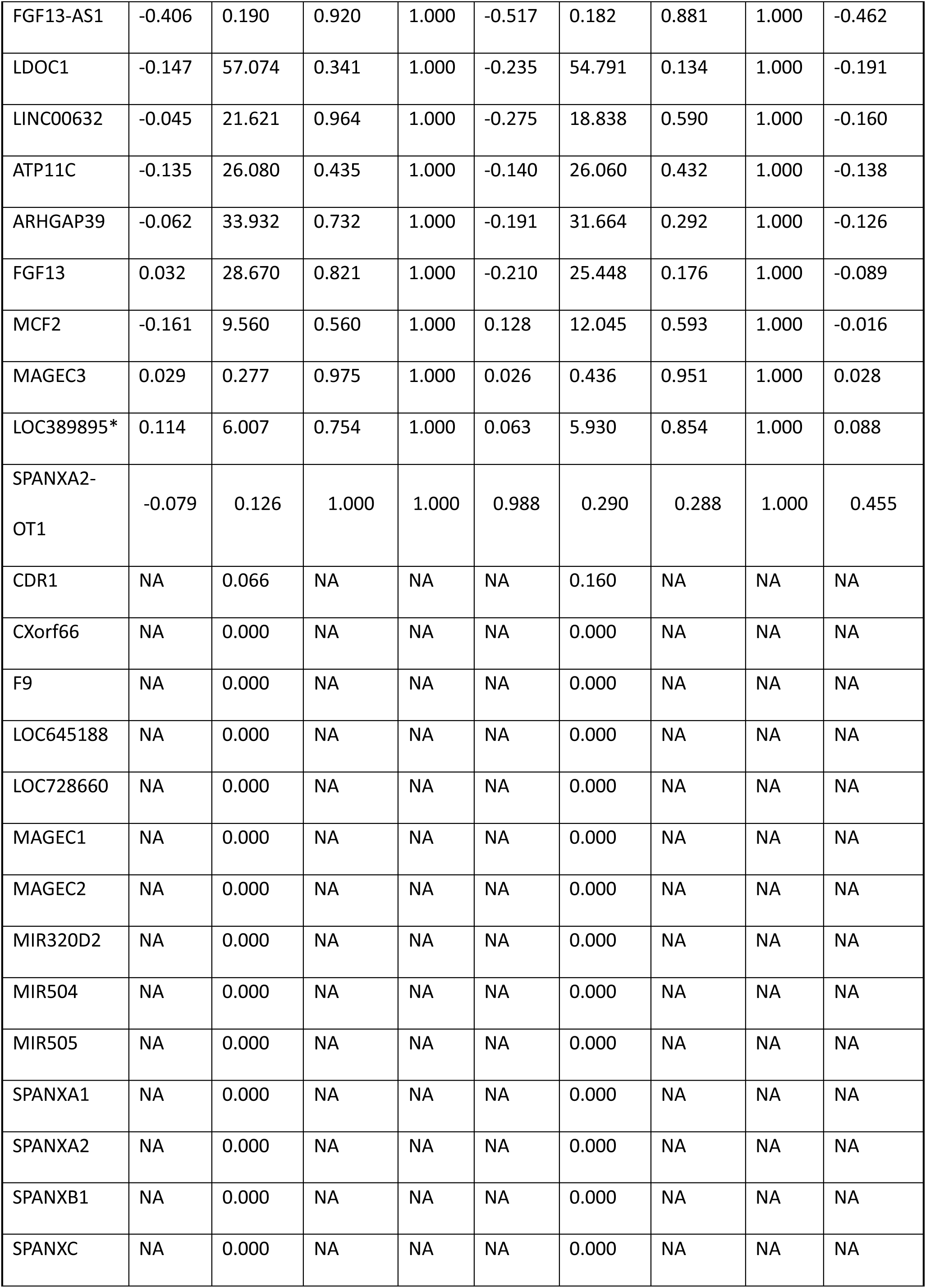
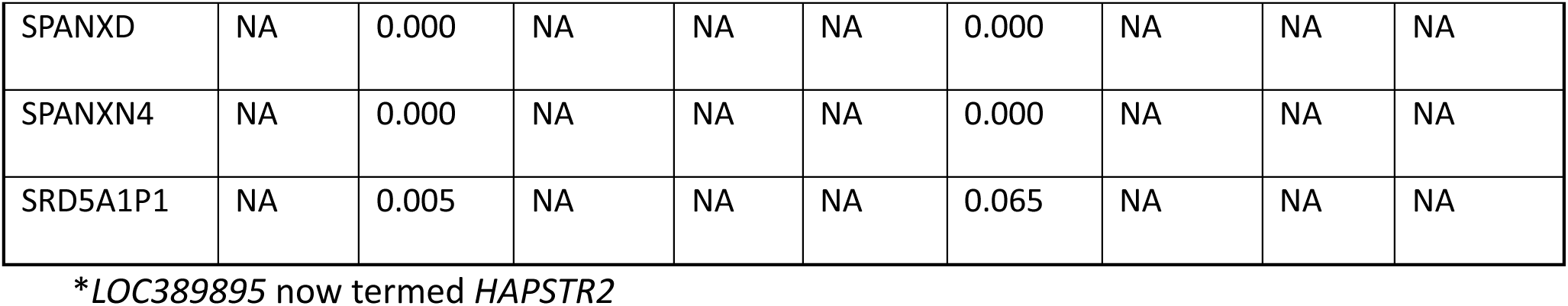
Exploratory differential expression of positional candidate genes in patient-derived sMN. Positional candidates include all Refseq curated genes located within the CMTX3 linkage interval and the partially duplicated *ARHGAP39* gene. RNA-Seq was performed on sMN mRNA isolated from 1 CMTX3 patient (P1: 2 iPSC clones, Clone A and Clone B) and 3 unrelated control lines. Differential expression was analysed using edgeR and comparing each patient clone separately against the control cohort (i.e. 1 vs 3). LogFC represents patient vs control. LogFC = Log(fold change); TPM = transcripts per kilobase million; FDR = false discovery rate; NA denotes genes excluded due to low/undetectable counts.

### Overexpression of the partially duplicated *ARHGAP39* is not detected in CMTX3 patients

To further evaluate potential overexpression of the partial *ARHGAP39* transcript in CMTX3, NanoString nCounter analysis was performed using probes targeting an exon localised within the CMTX3 SV and an exon outside the duplicated region. Both probes displayed similar levels of mRNA expression between patient and control samples across all developmental stages analysed, from iPSC to mature sMN (Figure 3).

**Figure 3:**
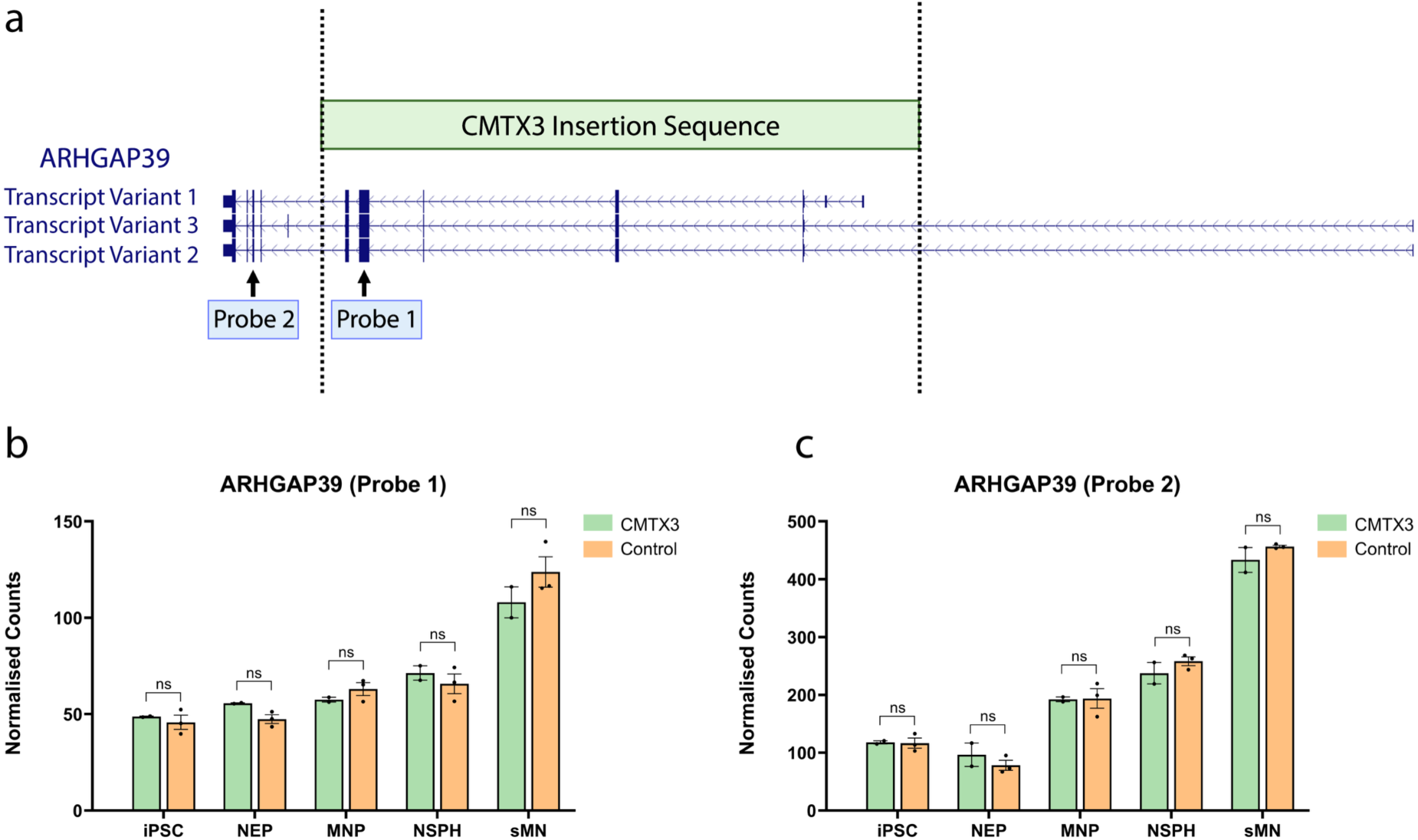
ARHGAP39 expression across iPSC to sMN differentiation. (A) Schematic of ARHGAP39 RefSeq curated transcript structure (UCSC Genome Browser, GRCh38) and the position of NanoString nCounter probes relative to the CMTX3 insertion sequence (green box). Probe 1 targets an exon within the partially duplicated ARHGAP39 segment contained within the CMTX3 insertion. Probe 2 targets an exon outside the duplicated region. Created with BioRender. (B-C) Normalised NanoString nCounter counts for ARHGAP39 measured using Probe 1 (B) and Probe 2 (C) in CMTX3 patient-derived (n = 2) and control (n = 3) lines across five stages of sMN differentiation. Data are shown as mean ± SEM. iPSC; induced pluripotent stem cells, NEP; caudal neuroepithelial progenitors, MNP; motor neuron progenitors, NSPH; neurospheres, sMN; spinal motor neurons.

### *SOX3* is spatiotemporally dysregulated in CMTX3 sMN differentiation

In parallel with the global transcriptomic analysis by RNA-Seq, the NanoString nCounter platform was used to conduct a targeted assessment of local gene regulation within the CMTX3 linkage interval. Custom probes were designed to measure the expression levels of 21 positional candidate genes located within ∼3 Mb either side of the CMTX3 SV (Supplementary Table 3) and expression was profiled at five stages of sMN differentiation (Figure 4A-E). At the iPSC stage, 12/21 candidate genes showed no detectable expression. Of the genes expressed, *SOX3* was the only transcript that differed significantly between CMTX3 and control iPSC (*p* = 0.0012), with patient-derived lines showing an approximately 5.7-fold reduction (log2(Ratio) = −2.51) (Figure 4A). At the caudal NEP stage, no candidate gene reached significance, however, *SOX3* again showed a trend towards reduced expression in CMTX3 samples (log2(Ratio) = −1.75, p = 0.157) (Figure 4B). Across the subsequent stages of differentiation (MNP, neurospheres and mature sMN) no positional candidate genes, including *SOX3,* were differentially expressed between CMTX3 and control lines (Figure 4C-E). Together, these results demonstrate spatiotemporally restricted reduction in *SOX3* expression that is most prominent at the iPSC stage. This pattern prioritises *SOX3* as a leading candidate for functional follow-up to define its potential pathogenic role in CMTX3. Quantification of *SOX3* expression dynamics at different timepoints of sMN differentiation is shown in Figure 5.

**Figure 4:**
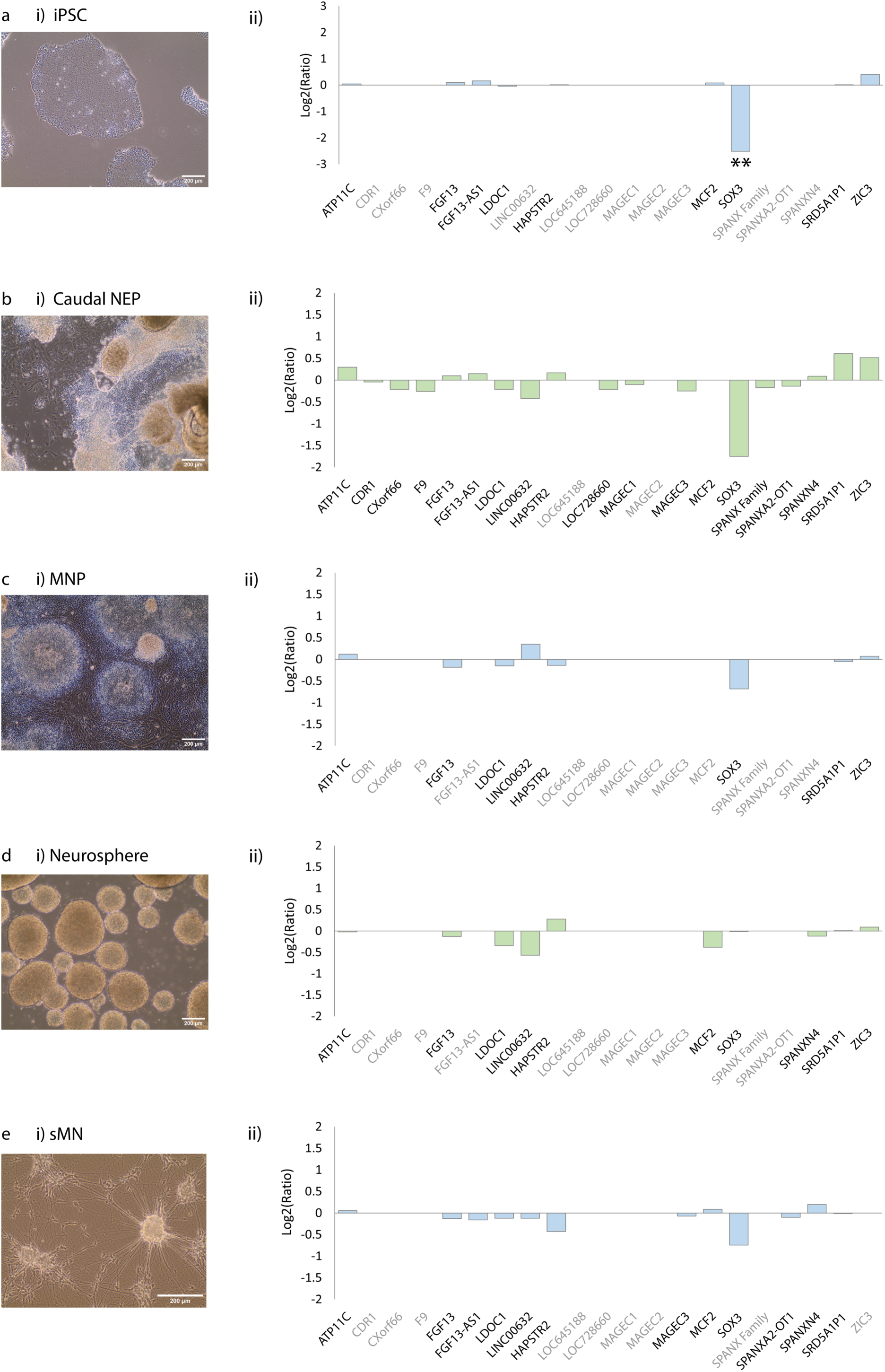
Relative expression of positional candidate genes throughout various stages of iPSC-to-sMN differentiation. RNA was isolated from CMTX3 (n=2) and control (n=3) samples at 5 discrete timepoints throughout the sMN differentiation process; (A) iPSC; (B) caudal NEP (differentiation day 6), (C) MNP (differentiation day 12), (D) neurospheres (differentiation day 18), and (E) sMN (differentiation day 28). Candidate gene expression was analysed using a custom NanoString nCounter panel. (i) Brightfield microscope images at each timepoint (scalebar = 200 μm). (ii) Relative expression of positional candidate genes, expressed as the log2(Ratio) of CMTX3 vs control. Candidate genes with undetectable expression levels in all samples are denoted in grey font. Statistical analysis was performed using nSolver Analysis software. **p < 0.01.

**Figure 5:**
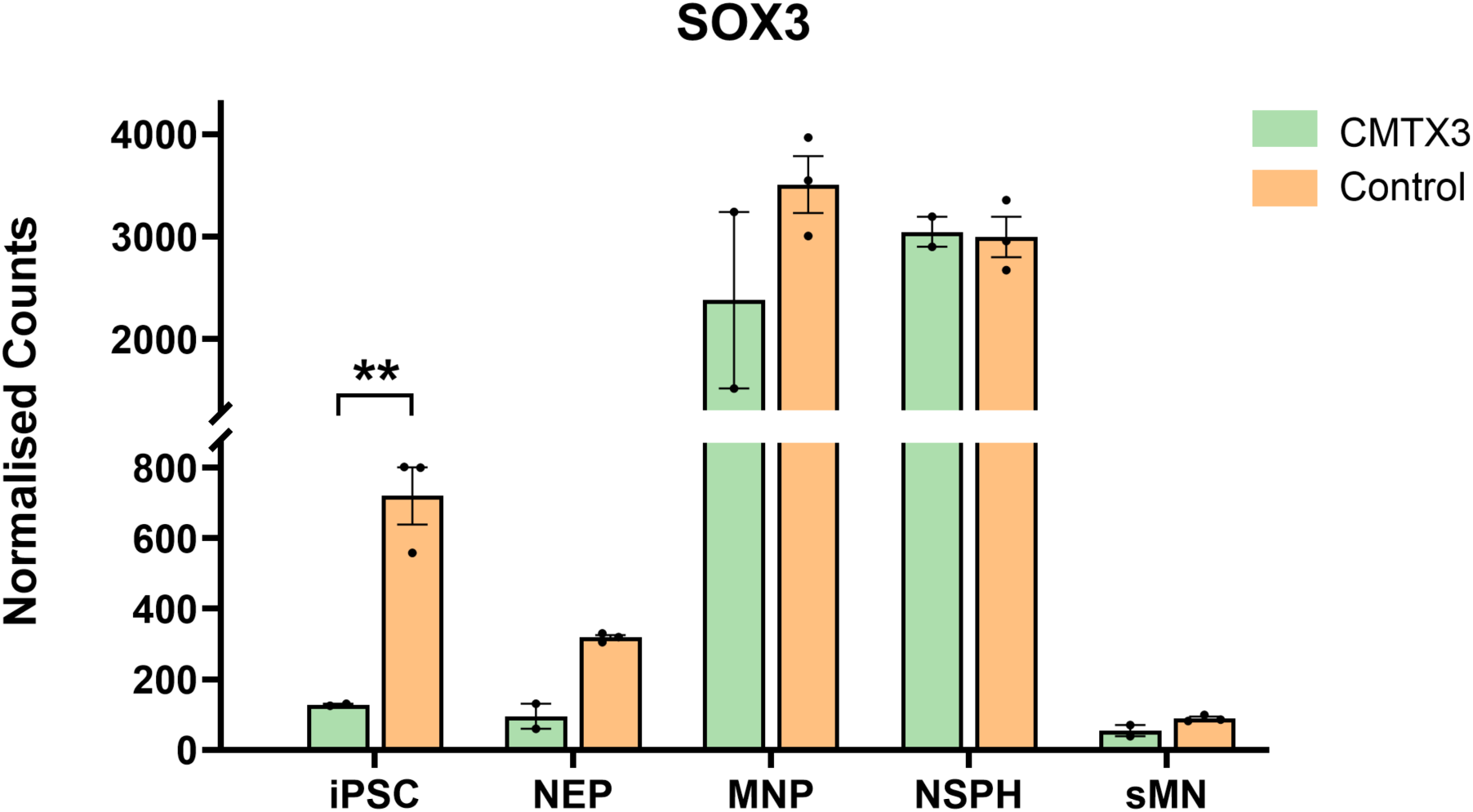
*SOX3* expression during iPSC-to-sMN differentiation in CMTX3. Nanostring nCounter counts for *SOX3* in CMTX3 patient-derived lines (n=2) and controls (n=3) across 5 stages of differentiation: iPSC, NEP, MNP, NSPH, and sMN. Bars show mean ± SEM. Differential expression analysis performed for each discrete timepoint using nSolver Analysis Software. ** *p* < 0.01.

Given that SOX3 functions as a transcription factor within the developing nervous system^27^, RNA-Seq was performed on RNA isolated from patient (n = 2) and control (n = 3) caudal NEP cultures to further explore the transcriptomic consequences of CMTX3. Quality control metrics are available in Supplementary Table 15. Differential gene expression analyses with DESeq2^20^ and edgeR^19^ identified 14 and 16 DEG, respectively. *SOX3* was one of the 14 DEG detected by DESeq2, and the only DEG located on chromosome X within the linkage region (Table 2A). CMTX3 NEP display a ∼3.89-fold (log2(FC) = −1.96) reduction in *SOX3* mRNA relative to controls (*p*-adj = 0.0015), consistent with the trend observed in the NanoString nCounter analysis. In contrast, none of the 16 DEG detected by edgeR map to chromosome X (Table 2B). Although *SOX3* did not reach significance with edgeR, the result narrowly missed the threshold (log2(FC)= −1.966, FDR = 0.0506), suggesting a biologically meaningful reduction. For completeness, NEP RNA-Seq datasets were also screened for the presence of novel fusion/splice/transcript variants involving CMTX3 genomic regions of interest as in the preliminary sMN RNA-Seq experiment. No CMTX3-specific features of interest were identified.

**Table 2:** Differentially expressed genes in caudal neuroepithelial progenitors from CMTX3 versus control lines. RNA-Seq was performed on poly(A) selected RNA isolated from caudal NEPs (differentiation day 6) derived from CMTX3 (n=2) and control (n=3) lines. (A) Differentially expressed gene results from DESeq2 with Log2FoldChange calculated using ‘control’ as the reference group and genes called significant at *p*-adjusted < 0.1 (default threshold). Genes are ranked by adjusted *p*-value. The CMTX3 positional candidate gene, *SOX3*, is highlighted in green. (B) Differentially expressed gene results from edgeR, with Log2FoldChange calculated using control as the reference group, and significance defined as FDR (false discovery rate) < 0.05. Genes are ranked by FDR.

| Gene | log2FoldChange | Adjusted *p*-value | Chr |
| --- | --- | --- | --- |
| LINC01139 | 3.671 | 1.30861E-11 | 1 |
| EIF2S3B | 3.781 | 1.58463E-06 | 12 |
| TMX2-CTNND1 | 3.398 | 9.82779E-06 | 11 |
| PLOD2 | -1.151 | 1.25649E-05 | 3 |
| SOX3 | -1.962 | 0.00149 | X |
| REC8 | -1.948 | 0.00303 | 14 |
| TRNY | -1.127 | 0.00367 | MT |
| AHRR | -9.207 | 0.01918 | 5 |
| TMEM132C | -1.343 | 0.01954 | 12 |
| PLEKHA2 | -1.277 | 0.02150 | 8 |
| TRNS1 | -1.137 | 0.03343 | MT |
| LOC112268219 | -7.415 | 0.03343 | 1 |
| MIR600HG | -4.269 | 0.03622 | 9 |
| CASKIN1 | -0.910 | 0.06108 | 16 |

**Table 2: Differentially expressed genes in caudal neuroepithelial progenitors from CMTX3 versus control lines.**
| Gene | log2FoldChange | FDR | Chr |
| --- | --- | --- | --- |
| LINC01139 | 3.660 | 2.33E-06 | 1 |
| TEKT4P2 | 11.275 | 1.33E-05 | 21 |
| EIF2S3B | 3.765 | 5.94E-05 | 12 |
| HLA-DRB1 | 7.946 | 9.09E-05 | 6 |
| TMX2-CTNND1 | 3.392 | 0.00043 | 11 |
| LOC112268219 | -7.960 | 0.00064 | 1 |
| PLOD2 | -1.156 | 0.00328 | 3 |
| BORCS7-ASMT | -7.637 | 0.00594 | 10 |
| REC8 | -1.954 | 0.00594 | 14 |
| TMX2P1 | 6.849 | 0.01027 | 9 |
| MRPL23-AS1 | -7.212 | 0.01368 | 11 |
| CCDC38 | 6.805 | 0.01416 | 12 |
| TMEM132C | -1.348 | 0.02414 | 12 |
| TRNY | -1.132 | 0.02414 | MT |
| TDGF1 | 6.514 | 0.03495 | 3 |
| RPL10P9 | 6.588 | 0.03839 | 5 |

### Proteomics confirms downregulation of SOX3 in CMTX3 neuroepithelial cells

To further validate the temporal dysregulation of *SOX3* observed at the mRNA level, iPSC from CMTX3 patients (n = 3) and controls (n = 3) were differentiated into caudal NEP then harvested for protein extraction for TMT LC-MS/MS analysis. Following TMT fragment quantification, removal of duplicate features and filtering to retain proteins identified in all samples from at least one condition, 8000 proteins were quantified per sample (Supplementary Figure 2). Only three proteins were encoded by genes residing within the CMTX3 linkage interval (ATP11C, SOX3, ARHGAP39; Supplementary Figure 3). Across the full proteome, 23 proteins were significantly altered between CMTX3 and control NEPs (*p.adj* < 0.05 Figure 6A; Supplementary Table 16). SOX3 was the only differentially abundant protein encoded within the CMTX3 linkage region (*p.adj* = 0.0252) and was reduced in CMTX3 NEPs relative to controls (Log2FC = −0.442, Figure 6B), concordant with the RNA-Seq and NanoString transcriptomic findings.

**Figure 6.**
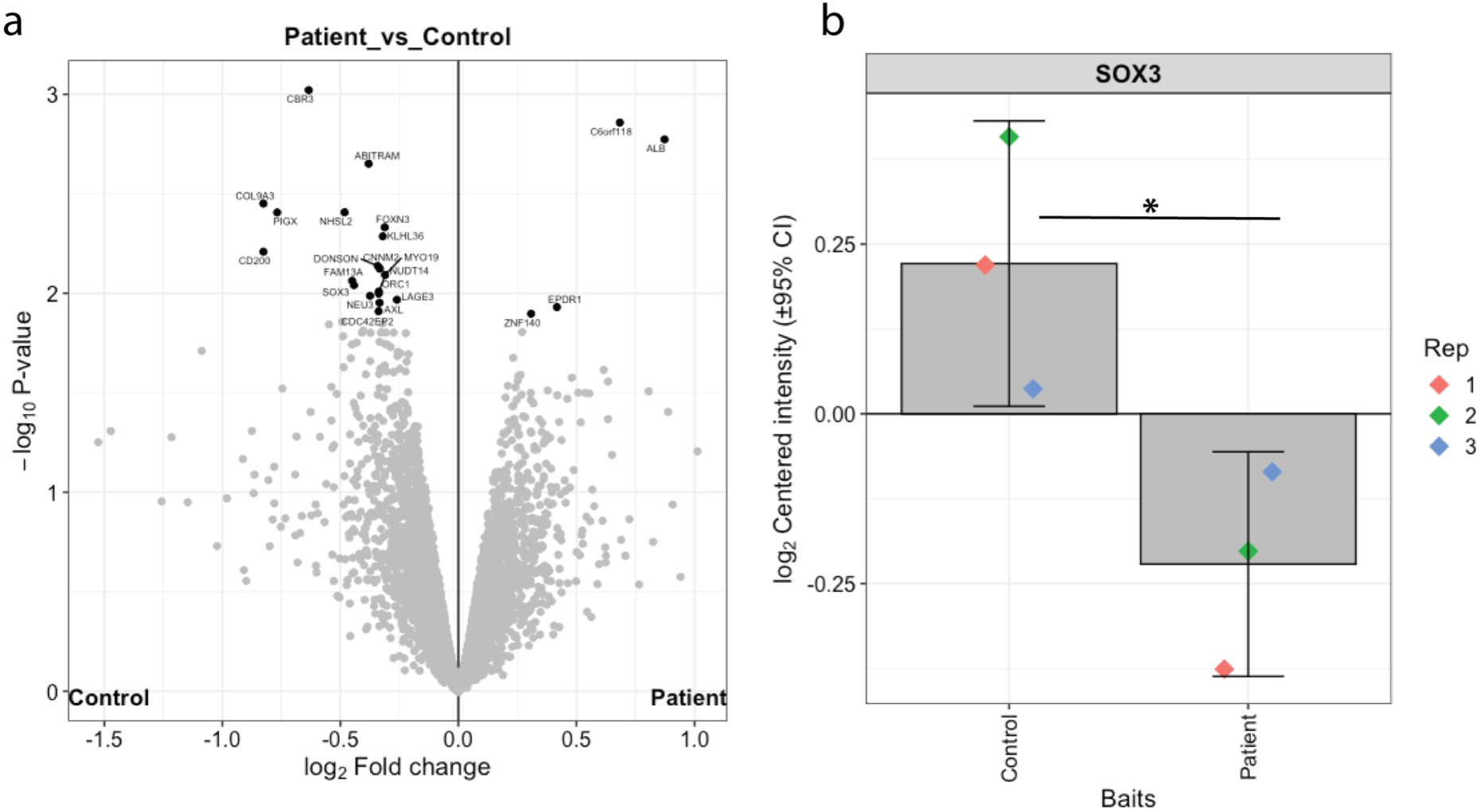
Proteomic profiling identifies reduced SOX3 abundance in CMTX3 patient-derived samples. (A) Volcano plot showing differential protein abundance between CMTX3 patient (n=3) and control (n=3) samples. The x-axis indicates log2 fold change (patient vs control) and the y-axis shows −log10 p-value. Proteins meeting the significance threshold are highlighted (black dots) and selected proteins are annotated. (B) Quantification of SOX3 protein abundance in control and CMTX3 patient samples, plotted as log2 centred intensity with 95% confidence intervals. Coloured points denote individual biological replicates (Rep 1–3). Differential expression was determined using DEP^25^. The asterisk (*) denotes a significant difference between groups (*adj. p* < 0.05).

## Discussion

Detecting complex pathogenic mutations has been greatly enhanced by modern advances in genomic sequencing technologies. The challenge now lies in interpreting their functional consequences. This is exemplified by CMTX3, in which the causative SV lies in non-coding DNA and does not disrupt any of the >130 genes^28^ known to cause CMT or related neuropathies. Establishing a mechanistic link between the CMTX3 associated genotype and clinical phenotype has been challenging.

In this study, we generated the first neuronal model of CMTX3, enabling the opportunity to investigate disease mechanism in relevant cell types that retain the disease associated SV. This is critical because the two leading hypotheses for CMTX3 pathogenesis (local gene dysregulation, or generation of a novel transcript) are transcription-based and therefore likely to be directly influenced by spatiotemporal context. Therefore, we evaluated both hypotheses in parallel, using two independent but complementary transcriptome platforms: short read RNA-Seq and NanoString nCounter.

Across all stages assessed during iPSC to sMN differentiation, neither the RNA-Seq nor NanoString nCounter provided evidence that the partially duplicated *ARHGAP39* region generates a novel fusion transcript or spliced isoform in CMTX3, and no *ARHGAP39* dosage effects were observed. Together, these data argue strongly against *ARHGAP39* as the causative gene for CMTX3. In contrast, both platforms independently demonstrated downregulation of the positional candidate, *SOX3*, in patient-derived cells confined to early developmental stages (iPSC and caudal NEP) and not observed at later stages of sMN differentiation. This patient-specific reduction of *SOX3* within caudal NEP samples was corroborated at the protein level by quantitative proteomics, supporting a model in which the CMTX3 SV perturbs the spatiotemporal regulation of *SOX3* during iPSC to sMN differentiation.

Notably, complex SVs at the Xq27.1 hotspot have previously been associated with local gene dysregulation, including XX-male sex reversal (*SOX3*)^29^ and X-linked congenital generalised hypertrichosis (*FGF13*)^30^, consistent with a shared regulatory mechanism at this locus. More recently, a distinct complex SV at the Xq27.1 hotspot was reported in an individual with clinical features overlapping CMTX3 neuropathy, accompanied by a tethered spinal cord, ophthalmological symptoms and neuropsychiatric features^31^. Although the mechanism was not resolved, these observations raise the possibility of convergent gene dysregulation^31^ and motivate testing whether *SOX3* dysregulation is also present in cells harbouring this alternative genomic rearrangement.

*FGF13* was previously proposed as a high-priority candidate gene based on its dysregulation in CMTX3 patient-derived lymphoblasts^5^. However, in the current study no consistent differences in *FGF13* expression were observed between patient and control cells at any stage of differentiation from iPSC to mature sMN. Among the positional candidates assessed, *SOX3* was the only gene to show reproducible CMTX3-associated expression changes. Collectively, these findings prioritise *SOX3* as the leading candidate gene underlying CMTX3 neuropathy.

*SOX3* is the closest gene to the CMTX3 structural variant, located ∼ 82 kb telomeric to the distal insertion breakpoint. It encodes the SOX3 transcription factor, a member of the SRY-related HMG box (SOX) family of transcription factors that play a role in embryonic development^32,33^. Together with SOX1 and SOX2, SOX3 forms the SOXB1 sub-family which is highly expressed in the developing vertebrate nervous system^34–37^ with a pivotal role in neural fate acquisition^27,36,38^. SoxB1 factors maintain neural progenitors in an undifferentiated state, thereby preventing premature terminal differentiation^39^. Beyond its well-established role in neurogenesis, SOX3 also contributes to gliogenesis. It has been reported to prevent premature differentiation of astrocytes in the developing spinal cord^40^, and conditional Sox3 deletion in mice impairs terminal differentiation of oligodendrocytes in the developing spinal cord^41^.

*SOX3* mutations are associated with a broad range of phenotypes, but *SOX3* has not been linked to CMT or related peripheral neuropathies. Pathogenic *SOX3* variants, including genomic duplications^42,43^, polyalanine tract expansions^42,44–46^, a polyalanine tract deletion^45^ and missense mutations^47,48^, most commonly present with variable hypopituitarism phenotypes, underscoring a key role for SOX3 in hypothalamo-pituitary development^42^. Clinical expressivity of hypopituitarism is highly variable, even among individuals carrying the same variant^48^, and is frequently accompanied by additional features including intellectual disability^43,44,47^, neural tube defects^43^, structural brain abnormalities^42,43,48^, and facial anomalies^44,47^ among others^47,48^.

Multiple duplications at the *SOX3* locus have been reported in patients with 46, XX disorders of sexual development (DSD) (SRY-negative), including XX-male sex reversal and XX-ovotesticular DSD^49–57^. Importantly, genomic rearrangements which leave the *SOX3* coding sequence intact have also been associated with 46, XX DSD. This includes a deletion immediately upstream of *SOX3*^50^, copy number gains occurring ∼ 566 kb upstream of *SOX3*^58^, and an inter-chromosomal insertion from chromosome 1 which is ∼ 82 kb distal to this gene^29^. These variants are believed to cause altered cis-regulatory architecture^58^ and potential ectopic *SOX3* expression during gonadal development^29,50^. Notably, the interchromosomal insertion reported by Haines et al. (2015) occurs within the quasi-palindromic sequence at Xq27.1^29^, a recognised hotspot for pathogenic SV, including the CMTX3 interchromosomal insertion (reviewed in Boyling et al. (2022)^7^). In summary, given the absence of established clinical associations between *SOX3* and peripheral neuropathy, further work is required to determine whether the *SOX3* dysregulation observed in CMTX3 is causally linked to the neuropathy phenotype.

*SOX3* is a key developmental transcription factor and was selectively dysregulated in early-stage developmental cell types derived from CMTX3 patients. This pattern raises the possibility that CMTX3 may arise, at least in part, from perturbed peripheral nerve development, consistent with its early-onset clinical presentation^59^. Developmental contributions have also been implicated in other forms of CMT. For example, perturbed Schwann cell differentiation was observed in both a rodent^60^ and iPSC^61^ model of CMT1A, supporting the concept that impaired developmental programmes may contribute to disease pathogenesis^61^.

Given ongoing debate as to whether CMTX3 is best classified as an axonal^62^, intermediate^4,5^ or demyelinating^59^ neuropathy, future studies should also assess candidate gene expression and regulation during Schwann cell differentiation. Although SOX3 emerges as a strong candidate based on its spatially and temporally restricted dysregulation, the current data do not demonstrate that altering SOX3 levels drives neuronal or glial phenotypes relevant to CMTX3 (e.g. axonal outgrowth, survival, myelination, electrophysiology). Establishing causality will require functional perturbation studies such as restoring SOX3 in patient cells, recapitulating SOX3 downregulation in control cells and developing in vivo models that reproduce the CMTX3-like pattern of SOX3 dysregulation. In this context, an auxin-inducible degron system could be particularly informative, providing a tuneable and reversible approach to modulate SOX3 protein levels across defined developmental stages^63^.

A limitation of the present study was the inability to assess the expression of miRNA genes within the CMTX3 linkage region (*MIR504, MIR505, MIR320D2*), as the experimental design was optimised for robust quantification of coding and long non-coding transcripts. Therefore, we cannot exclude the possibility that one or more miRNAs are dysregulated by the complex insertion. A further limitation is the small number of iPSC lines analysed and the relatedness of the patient-derived lines. All reported CMTX3 patients harbouring the ∼78 kb insertion of 8q24.3 originate from three pedigrees sharing a common ancestor in which the mutation first arose^64^. As such, the observed transcriptional differences could be influenced by family-specific background variation. Expanding to larger cohorts and/or incorporating isogenic controls would strengthen the link between CMTX3 and *SOX3* dysregulation. Although multiple differentiation timepoints were sampled, they may not capture the earliest developmental windows in which SOX3 exerts its effects on neural differentiation. Finally, bulk RNA-seq and NanoString measure average expression across heterogeneous cell populations, which may mask subtle or subpopulation–specific changes that could be resolved using single-cell and/or spatial transcriptomic approaches. Coupling these advanced technologies with PNS organoid models^65^ of CMTX3 could further explore the impact of *SOX3* dysregulation on PNS development.

## Conclusion

Overall, this study highlights the challenge of interpreting non-coding structural variants associated with inherited neuropathies. Using the first neuronal model of CMTX3, RNA-seq and NanoString supported by quantitative proteomics converged on a reproducible, patient-specific reduction of SOX3 restricted to early developmental stages and consistent with disrupted spatiotemporal regulation driven by the CMTX3 insertion. These findings position *SOX3* as a novel, leading candidate gene for CMTX3 and support a model in which Xq27.1 hotspot rearrangements remodel local regulatory architecture. Collectively, these data represent a significant step toward elucidating the pathomechanism of CMTX3 and further support a model in which altered gene regulation may underly this neuropathy.

## Acknowledgements

The authors gratefully acknowledge the CMT families who have participated in this research. This work was supported by National Health and Medical Research Council Project Grant (APP11868687) warded to M.L.K., MRFF Genomics Health Futures Mission Grant (APP2007681) awarded to M.L.K and S.V., a CMT Association Australia grant awarded to M.L.K. and A.B., a National Health and Medical Research Council Investigator Grant (GNT2008066) awarded to J.J.D., and an Australian Government Research Training Program (RTP) Scholarship awarded to A.B. This article is based on work included in A.B.’s PhD thesis (The University of Sydney).

Supplementary information accompanies the manuscript.

