## Supplementary Material for "The Charcot-Marie-Tooth Neuropathy (CMTX3) Complex Structural Variation Causes Differential SOX3 Spatiotemporal Expression"

**Supplementary Table 1:** Summary of iPSC lines used in the present study

| **Cell Line ID** | **Donor Sex** | **Donor Race** | **iPSC Source** | **Experiments Utilised** |
| --- | --- | --- | --- | --- |
| CMTX3 Patient 1 (P1) | Male | Caucasian | Fibroblasts reprogrammed by FUJIFILM Cellular Dynamics (Wisconsin, USA) | Preliminary RNA-Seq screen on sMN  NanoString Analyses  NEP RNA-Seq  Proteomics |
| CMTX3 Patient 2 (P2) | Male | Caucasian | Fibroblasts reprogrammed by Stem Cell and Organoid Facility (Children’s Medical Research Institute, Westmead, Australia) | NanoString Analyses  NEP RNA-Seq  NEP Proteomics |
| CMTX3 Patient 3 (P3) | Male | Caucasian |  | NEP Proteomics |
| Control 1 (C1) | Male | Caucasian | iPSC line purchased from  FUJIFILM Cellular Dynamics (Wisconsin, USA) #CW60014BB1 | Preliminary RNA-Seq screen on sMN  NanoString Analyses  NEP RNA-Seq  NEP Proteomics |
| Control 2 (C2) | Male | Caucasian | iPSC line purchased from  FUJIFILM Cellular Dynamics (Wisconsin, USA) #CW60304EE1 |  |
| Control 3 (C3) | Male | Caucasian | Fibroblasts reprogrammed by FUJIFILM Cellular Dynamics (Wisconsin, USA) |  |

**Supplementary Table 2:** Taqman Gene Expression Assays used in study

| **Gene ID** | **Taqman Assay ID** |
| --- | --- |
| *LIN28A* | Hs00702808_s1 |
| *NANOG* | Hs02387400_g1 |
| *NODAL* | Hs00415443_m1 |
| *FOXD3* | Hs00255287_s1 |
| *CDH1* | Hs00170423_m1 |
| *TDGF1* | Hs02339499_g1 |
| *GAPDH* | Hs99999905_m1 |

**Supplementary Table 3:** Genes targeted by the custom NanoString nCounter Elements TagSet panel.

| **Gene** **Target** | **Class** | **Comment** | **Isoform coverage (# hit/ # total)** |
| --- | --- | --- | --- |
| ARHGAP39* | Target | Partially duplicated in CMTX3 | 6/6 |
| ARHGAP39* | Target | Partially duplicated in CMTX3 | 6/6 |
| ATP11C | Target | CMTX3 positional candidate | 18/18 |
| CDR1 | Target | CMTX3 positional candidate | 1/1 |
| CHAT | Target | Differentiation Marker | 7/7 |
| CXorf66 | Target | CMTX3 positional candidate | 1/1 |
| EGR2 | Target | Differentiation Marker | 6/6 |
| F9 | Target | CMTX3 positional candidate | 3/3 |
| FGF13 | Target | CMTX3 positional candidate | 8/8 |
| FGF13-AS1 | Target | CMTX3 positional candidate | 1/1 |
| GAP43 | Target | Differentiation Marker | 3/3 |
| ISL1 | Target | Differentiation Marker | 2/2 |
| LDOC1 | Target | CMTX3 positional candidate | 1/1 |
| LINC00632 | Target | CMTX3 positional candidate | 1/3 |
| LOC389895 (HAPSTR2) | Target | CMTX3 positional candidate | 1/1 |
| LOC645188 | Target | CMTX3 positional candidate | 1/1 |
| LOC728660 | Target | CMTX3 positional candidate | 1/1 |
| MAGEC1 | Target | CMTX3 positional candidate | 2/2 |
| MAGEC2 | Target | CMTX3 positional candidate | 1/1 |
| MAGEC3 | Target | CMTX3 positional candidate | 4/4 |
| MCF2 | Target | CMTX3 positional candidate | 12/12 |
| MNX1 | Target | Differentiation Marker | 2/2 |
| MPZ | Target | Differentiation Marker | 3/3 |
| NES | Target | Differentiation Marker | 1/1 |
| NOTCH1 | Target | Differentiation Marker | 2/2 |
| OLIG2 | Target | Differentiation Marker | 2/2 |
| PMP22 | Target | Differentiation Marker | 9/9 |
| SOX2 | Target | Differentiation Marker | 1/1 |
| SOX3 | Target | CMTX3 positional candidate | 1/1 |
| SPANX family** | Target | CMTX3 positional candidate |  |
| SPANXA2-OT1 | Target | CMTX3 positional candidate | 1/1 |
| SPANXN4 | Target | CMTX3 positional candidate | 1/3 |
| SRD5A1P1 | Target | CMTX3 positional candidate | 1/1 |
| ZIC3 | Target | CMTX3 positional candidate | 2/2 |
| CLTC | Housekeeping |  | 2/2 |
| G6PD | Housekeeping |  | 3/3 |
| GUSB | Housekeeping |  | 16/16 |
| HPRT1 | Housekeeping |  | 1/1 |
| PGK1 | Housekeeping |  | 1/1 |
| POLR1B | Housekeeping |  | 10/10 |
| POLR2A | Housekeeping |  | 1/1 |
| RPL30 | Housekeeping |  | 1/1 |
| RPLP0 | Housekeeping |  | 2/2 |
| SDHA | Housekeeping |  | 8/8 |
| TBP | Housekeeping |  | 2/2 |
| TFRC | Housekeeping |  | 10/10 |
| TUBB | Housekeeping |  | 7/7 |
| YWHAZ | Housekeeping |  | 11/13 |

* Two probes for *ARHGAP39* were included in the custom panel; one probe targeting sequence contained within the CMTX3 insertion fragment, and one probe located outside of this region.

**High sequence homology between *SPANX* family gene members prohibited unique probe design. Instead, a singular probe was designed to target *SPANXA1, SPANXA2, SPANXB1, SPANXC* and *SPANXD* @ >95%.

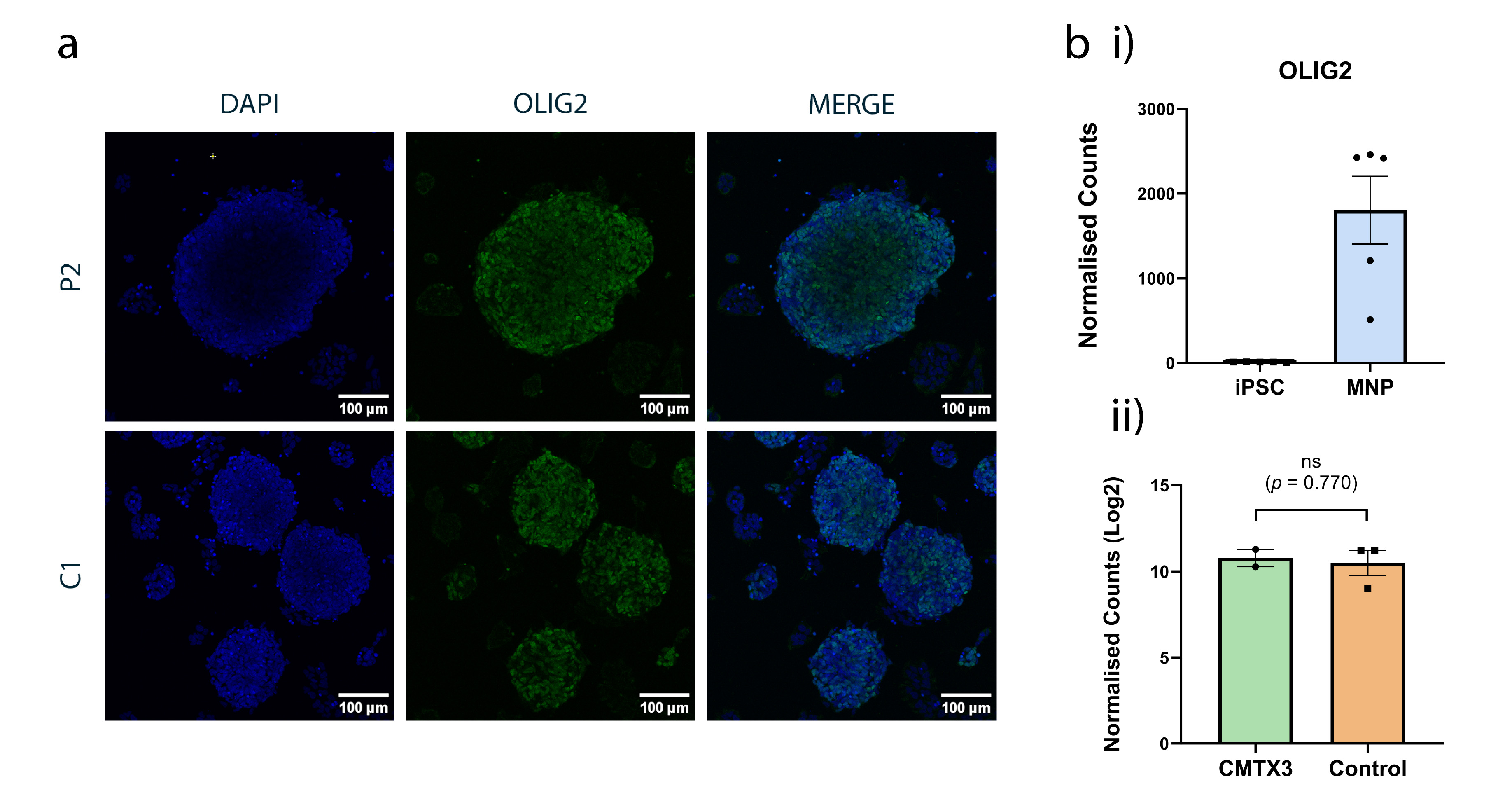

**Supplementary Figure 1: Generation of OLIG2-positive MNP.** (A) Representative immunofluorescence images showing OLIG2 (green) expression in iPSC-derived MNPs from a CMTX3 line (P2) and a control line (C1). MNP were stained 3-days after plating. Nuclei are counterstained with DAPI (blue); merged images are shown. (B) NanoString nCounter quantification of *OLIG2* mRNA (i) Normalised counts in iPSCs versus MNPs (n=5 lines per stage) demonstrating induction of *OLIG2* upon differentiation. (ii) Log2- transformed normalised counts comparing MNPs from CMTX3 (n=2) and control (n=3) lines; no significant difference was detected (t-test with Welch’s correction; ns, p=0.770). Bars represent mean ± SEM and points indicate individual cell lines. Data shown are from a single differentiation experiment.

****Supplementary Tables 4-15 are provided in an “Supplementary_Data_Tables.xlsx”**

**Supplementary Table 4:** Quality control metrics for sMN RNA-Seq libraries. GC = guanine-cytosine content; Q20 = ratio of bases with Phred quality score ≥ 20; Q30 = ratio of bases with Phred quality score ≥ 30.

**Supplementary Table 5:** Fusion transcripts with breakpoints on Chromosomes X and 8 predicted in CMTX3 patient-derived sMN. FusionCatcher predicted fusion events with breakpoints spanning chromosome X and 8 in CMTX3 sMNs (n = 1 individual; 2 iPSC clones). The other fusion detection tools (DeFuse and Arriba) did not identify any X-8 fusion in these samples. CMTX3 genomic regions of interest include the duplicated 8q24.3 segment (chr8:144,542,928-144,620,773) and regions surrounding the Xq27.1 breakpoint (chrX:140,420,783-140,420,784) (GRCh38).

**Supplementary Table 6:** Fusion transcripts, predicted by Arriba, on chromosome X from CMTX3 patient-derived sMN (n = 1 patient (2 clones)).

**Supplementary Table 7:** Fusion transcripts, predicted by deFuse, on chromosome X from CMTX3 patient-derived sMN (n = 1 patient (2 clones)).

**Supplementary Table 8:** Fusion transcripts, predicted by FusionCatcher, on chromosome X from CMTX3 patient-derived sMN (n = 1 patient (2 clones)).

**Supplementary Table 9:** Fusion transcripts, predicted by deFuse, on chromosome 8 from CMTX3 patient and control sMN (n = 1 patient (2 clones)). A fusion transcript involving *ARHGAP39*, which is partially duplicated in CMTX3, was identified but is found in controls as well as patient samples.

**Supplementary Table 10:** Fusion transcripts, predicted by Arriba, on chromosome 8 from CMTX3 patient-derived sMN (n = 1 patient (2 clones)).

**Supplementary Table 11:** Fusion transcripts, predicted by FusionCatcher, on chromosome 8 from CMTX3 patient-derived sMN (n = 1 patient (2 clones)). Due to the high number of predicted transcripts involving chromosome 8, only those with a breakpoint located within 3 Mb of the genomic region of interest (chr8:144,542,928 – 144,620,773) are shown.

**Supplementary Table 12:** Novel transcript variants on chromosomes 8 and X across CMTX3 and control-derived sMN (n = 1 patient (2 clones)). Transcripts were assembled with StringTie and classified with GffCompare. (A) Data from chromosome 8. (B) Data from chromosome X. Reads represented as TPM (transcripts per kilobase million). ‘Class code’ refers to the classification code assigned by the Gffcompare program.

**Supplementary Table 13:** Novel splice variants at genomic regions of interest on chromosomes X and 8 across CMTX3 and control-derived sMN (n = 1 patient (2 clones)). Transcripts were assembled with StringTie and classified with GffCompare. (A) Predicted splice variants located within 3 Mb either side of the breakpoint of the CMTX3 SV on Xq27.1. (B) Predicted splice variants falling within a genomic window extending 100 kb either side of the region of chromosome 8q24.3 that is duplicated in CMTX3 (chr8:144,542,928 – 144,620,773). Reads represented as TPM (transcripts per kilobase million). ‘Class code’ refers to the classification code assigned by the GffCompare program.

**Supplementary Table 14:** List of predicted DEG in CMTX3 sMN. DEG were identified using edgeR with FDR cutoff < 0.05. Each CMTX3 clone from P1 was individually compared to a cohort of 3 controls. Only those DEG identified in both clones, showing the same direction of logFC, are listed. Genes are listed alphabetically. logFC = log2(fold change); FDR = false discovery rate.

**Supplementary Table 15:** Quality control metrics for NEP RNA-Seq libraries. GC = guanine-cytosine content; Q20 = ratio of bases with Phred quality score ≥ 20; Q30 = ratio of bases with Phred quality score ≥ 30.

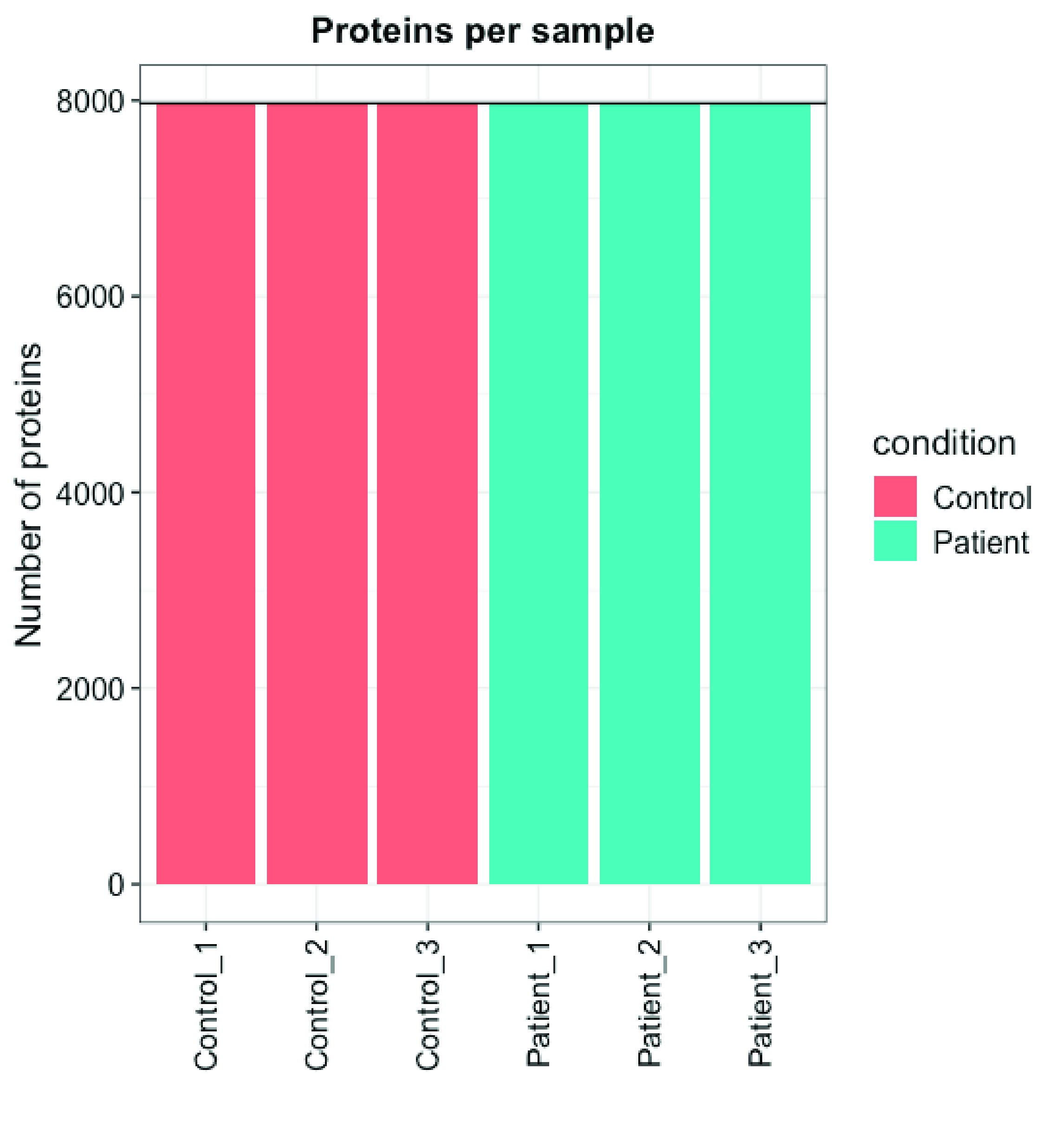

**Supplementary Figure 2:** QC of Protein Detection in CMTX3 and Control NEP samples. Bar graph representing the number of proteins detected in each sample by proteomics. 8000 proteins were detected in all samples

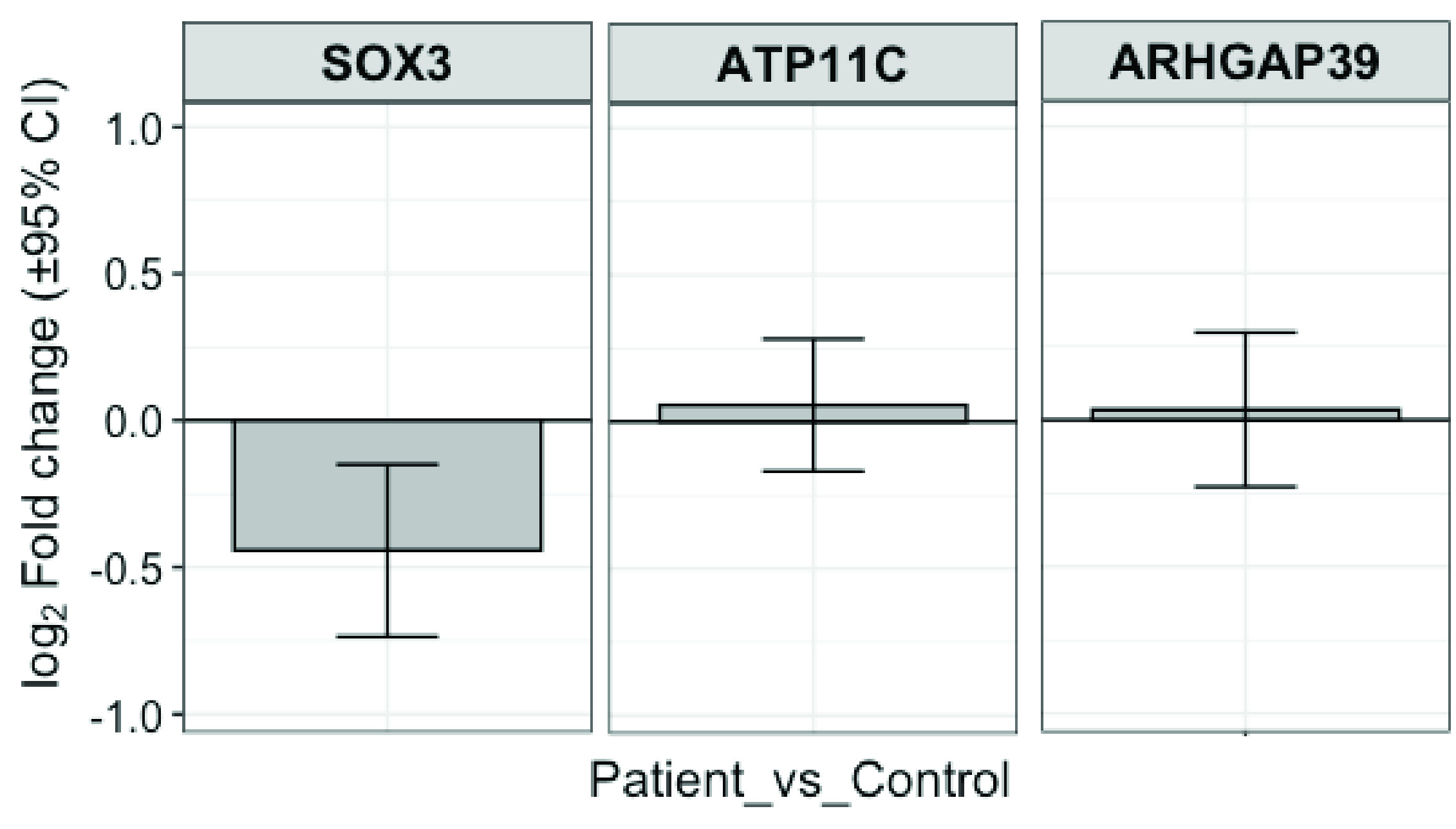

**Supplementary Figure 3:** Differential protein expression of genes within the CMTX3 linkage interval. Bar graphs showing Log2 fold change ratios (patient:control) of linkage interval proteins detected by proteomics. Error bars represent 95% Confidence Intervals.

| **name** | **ID** | **Ratio (P vs C)** | **P vs C (p.val)** | **P vs C (p adj)** |
| --- | --- | --- | --- | --- |
| CBR3 | O75828 | -0.634 | 0.00095248 | 2.86E-06 |
| C6orf118 | Q5T5N4 | 0.684 | 0.00138615 | 1.69E-05 |
| ALB | P02769 | 0.873 | 0.00168525 | 3.69E-05 |
| ABITRAM | Q9NX38 | -0.38 | 0.00223344 | 0.000137 |
| COL9A3 | Q14050 | -0.826 | 0.00353909 | 0.00101 |
| NHSL2 | Q5HYW2 | -0.482 | 0.00391493 | 0.00135 |
| PIGX | Q8TBF5 | -0.768 | 0.00392547 | 0.00136 |
| FOXN3 | O00409 | -0.312 | 0.00465878 | 0.0026 |
| KLHL36 | Q8N4N3 | -0.32 | 0.00517331 | 0.00367 |
| CD200 | P41217 | -0.826 | 0.00617925 | 0.00701 |
| DONSON | Q9NYP3 | -0.342 | 0.00729413 | 0.0118 |
| CNNM2 | Q9H8M5 | -0.333 | 0.00752575 | 0.0129 |
| MYO19 | Q96H55 | -0.31 | 0.00808595 | 0.0157 |
| FAM13A | O94988 | -0.45 | 0.00864722 | 0.0188 |
| SOX3 | P41225 | -0.442 | 0.00910565 | 0.0213 |
| NUDT14 | O95848 | -0.337 | 0.00979943 | 0.0252 |
| ORC1 | Q13415 | -0.337 | 0.01003712 | 0.0265 |
| NEU3 | Q9UQ49 | -0.375 | 0.01030227 | 0.0279 |
| LAGE3 | Q14657 | -0.26 | 0.01077812 | 0.031 |
| AXL | P30530 | -0.334 | 0.0111594 | 0.0334 |
| EPDR1 | Q9UM22 | 0.417 | 0.01173836 | 0.0376 |
| CDC42EP2 | O14613 | -0.338 | 0.01231291 | 0.0417 |
| ZNF140 | P52738 | 0.308 | 0.01265886 | 0.0442 |

**Supplementary Table 16:** Differentially expressed proteins between CMTX3 (n=3) and control (n=3) caudal NEP samples
